# Interaction between the assembly of the ribosomal subunits: Disruption of 40S ribosomal assembly causes accumulation of extra-ribosomal 60S ribosomal protein uL18/L5

**DOI:** 10.1101/756114

**Authors:** Nusrat Rahman, Md Shamsuzzaman, Lasse Lindahl

## Abstract

Inhibition of the synthesis of a ribosomal protein (r-protein) abrogates the assembly of its cognate subunit, while assembly of the other subunit continues. Ribosomal components that are not stably incorporated into ribosomal particles due to the disrupted assembly are rapidly degraded. The 60S protein uL18/L5 is an exception, because this protein accumulates extra-ribosomally during inhibition of 60S assembly. Since the r-proteins in each ribosomal subunit are essential only for formation of their own subunit, it would be predicted that accumulation of extra-ribosomal uL18/5 only occurs during restriction of 60S assembly, and not during abolition of 40S assembly. Contrary to this prediction, we report here that repression of 40S r-protein genes does in fact lead to accumulation of uL18/L5 does outside the ribosome due modified 60S assembly. Furthermore, the effect varies depending on which 40S ribosomal protein is repressed. We propose that disruption of early steps in the 40S subunit assembly changes the kinetics of 60S subunit assembly resulting in a buildup of extra-ribosomal uL18/L5, even though 60S formation continues. Finally, our results show that maintenance of the pool of extra-ribosomal uL18/L5 requires continual protein synthesis showing that extra-ribosomal protein is not stable, but is slowly “consumed” by incorporation into 60S subunits and/or turnover.

## Introduction

The ribosome biogenesis process is preserved throughout eukaryotic evolution, although the complexity has evolved from yeast to humans [1, 2]. It begins with RNA polymerase I transcription of a long 18S-5.8S-25S/28S precursor rRNA and RNA polymerase III transcription of precursor 5S rRNA [3–6]. The precursor transcripts are processed into the mature rRNA components concurrently with the incorporation of ribosomal proteins (r-proteins) into the emerging ribosomal subunits. Ribosomal proteins are translated in the cytoplasm and chaperoned into the nucle(o)lus where most of the ribosome formation takes place. Besides the synthesis of the components of the mature ribosomes, the construction of ribosomes also requires in excess of 250 ribosomal assembly factors, a number of which are important for the assembly of both the 40S and the 60S ribosomal subunits, while others are specific to the formation of one the ribosomal subunits [7].

Assembly of each subunit requires production of a full set of the r-proteins found in the mature subunit. Significant reduction of the production of just one r-protein or assembly factor prevents completion of the assembly process. If the perturbation is limited to protein(s) required for only one or the other subunit, only the assembly of the cognate subunit is abolished, while the assembly of the other subunit continues (e.g. [8, 9]). The abolishment of the assembly of a ribosomal subunit does not stop the synthesis of its r-proteins, but proteins that fail to become incorporated into stable ribosomal particles are rapidly eliminated by proteasomal turnover [8, 10, 11].

Nevertheless, one 60S protein, uL18, evades rapid degradation and accumulates in a complex with 5S rRNA outside of the ribosome during repression of the gene for the 60S r-protein uL5 [12], previously named L11 and L16 [13, 14]. Furthermore, distortion of cell fate in metazoans has been attributed to r-protein mediated regulation of factors controlling the growth [15, 16]. It is therefore important to understand the mechanisms for the build-up of extra-ribosomal r-protein pools. Since the r-proteins in each ribosomal subunit are essential only for the assembly of their cognate subunit, it would be expected that interruption of the assembly of one subunit only affects accumulation of extra-ribosomal r-proteins for that subunit. We tested this expectation by repressing several 40S r-protein genes and measuring the buildup of extra-ribosomal r-proteins.

Surprisingly, and in contrast to the prediction, extra-ribosomal uL18 accumulates when the synthesis of either 40S or 60S r-proteins is constrained. Moreover, the amount of extra-ribosomal uL18 accumulating depends on which 40S r-protein gene is repressed. We interpret these results to mean that disruption of the assembly of the 40S subunit can affect the kinetics of assembly of the 60S subunit. Furthermore, we show that buildup of extra-ribosomal uL18 does not require formation of the complex of uL18, uL5, 5S rRNA and the Rrs1 and Rpf2 assembly factors, which is an intermediate in the normal 60S subunit assembly.

## Materials and methods

### Nomenclature for r-proteins

We use the 2014 universal nomenclature [13]. In the figures, the classic protein names are also indicated after a slash.

### Strains and growth conditions

All strains are derived from BY4741. In each strain one r-protein gene encoding eS4, eS6, uS17, eS19, eS31, eL40, or eL43, was expressed exclusively from the *Gal1/10* promoter. These strains are referred to as P_gal_-xx where xx is the protein expressed from the galactose promoter (Table S1). In the experiment in Fig 1B described below, P_gal_-eL43 was transformed with a second plasmid carrying a gene for uL18-FLAG expressed from the constitutive RpS28 promoter (Philipp Milkereit, personal communication).

**Fig 1.**
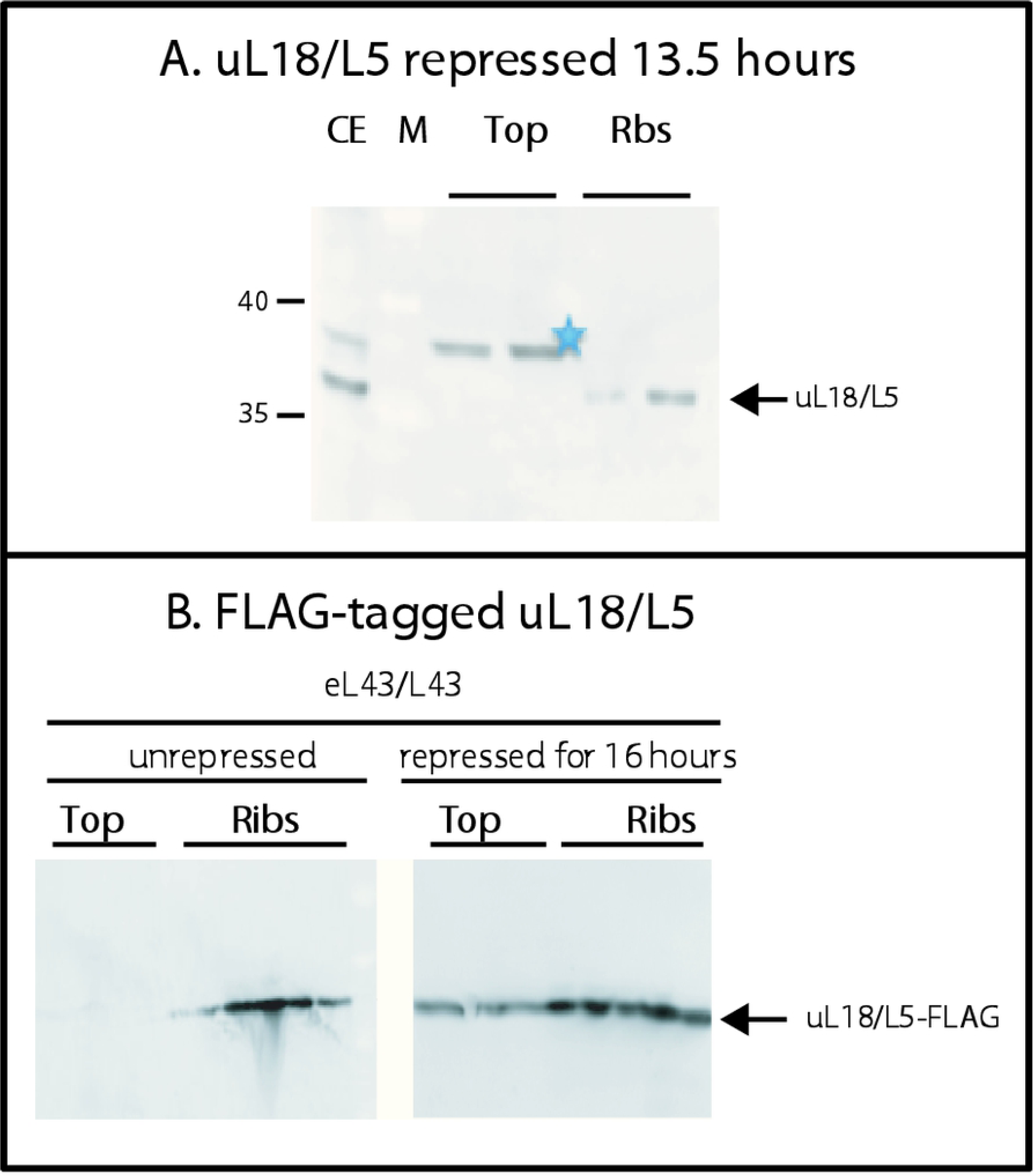
Analysis of the specificity of anti-uL18/L5. (A) P_gal_-uL18/L5 was grown in galactose medium and shifted to glucose medium. A lysate prepared after repression of uL18/L5 gene for 13.5 hours was fractionated on a sucrose gradient. Fractions from the top of the gradient and the 60S-80S ribosome peaks were analyzed by western blot stained with anti-uL18/L5. (B) P_gal_-eL43/L43 was transformed with a plasmid harboring a constitutively expressed gene for uL18/L5-FLAG. The culture was grown in galactose medium and shifted to glucose medium for 16 hours. Lysates prepared from cells before and after repressing uL43/L43 synthesis were fractionated on sucrose gradient and aliquots of fractions from the top of the gradient and the 60S-80S peaks were analyzed for content of FLAG-tagged protein by western blot.

Cells were grown at 30°C with shaking in YEP-galactose media. At OD_600_ of 1.0 (about 2×10^7^ cells per ml), the culture was shifted to YPD (glucose) media for 6-21 hours. All strains had a doubling time of 1.5-2.0 hours in galactose. After the shift, growth of the P_gal_-xx strains gradually decreased [9]. Cells were harvested at 8000 rpm for 10 minutes and washed once with 10 mL ice cold RNase free water and stored at - 20°C until further use. Procedures for lysis and sucrose gradient centrifugation were described previously [9].

### Western analysis and antisera

Western blots [9] were probed with rabbit polyclonal antisera prepared for our laboratory by Covance (Princeton, New Jersey, USA) using synthetic peptides with the sequence of 20–22 N-terminal amino acids of uS4, uL4, uL5, and uL18 as antigens. Monoclonal anti-FLAG antibody was purchased from Thermo-Fisher (catalogue number MA1-91878).

As described in Results, western blots probed anti-uL18 revealed a band co-migrating in electrophoresis with the ribosomal uL18 band in fractions close to the top of sucrose gradients loaded with extracts of specific strains shifted to glucose medium. To determine if this band indeed represents uL18 we analyzed P_gal_-uL18 13.5 hours after shifting the culture to glucose medium. In this experiment, we saw no band co-migrating with the ribosomal uL18 band at the top of the gradient, consistent with the fact the uL18 synthesis was abolished and confirming that the band at the top of the gradient, which co-migrates in gel electrophoresis and is stained with our anti-uL18, in fact is uL18 (Fig 1A). However, our anti-uL18 also reacted with a second band at the top of the sucrose gradients (marked with asterisk in all figures). To determine if this band is related to uL18, we transformed P_gal_-eL43 with a second plasmid constitutively expressing FLAG-tagged uL18 (see above) in addition to the native uL18 chromosomal gene. After the shift from galactose to glucose medium, a single band of FLAG-tagged uL18, comigrating with the FLAG-uL18 band in the ribosomal fractions, appeared at the top of the gradient (Fig 1B), but no band corresponding asterisked band in blots stained with anti-uL18 was seen. Furthermore, the asterisked band was present after repressing uL18 synthesis in Pgal-uL18 (Fig 1A). Based on the experiments in Fig 1, we conclude that the band marked with asterisk in the blots stained with anti-uL18 is not related to uL18, but must represent an unspecific protein that cross-reacts with our uL18 antiserum. This is supported by the presence of the asterisked band after shifting the parent strain to glucose medium (Fig 2C).

**Fig 2.**
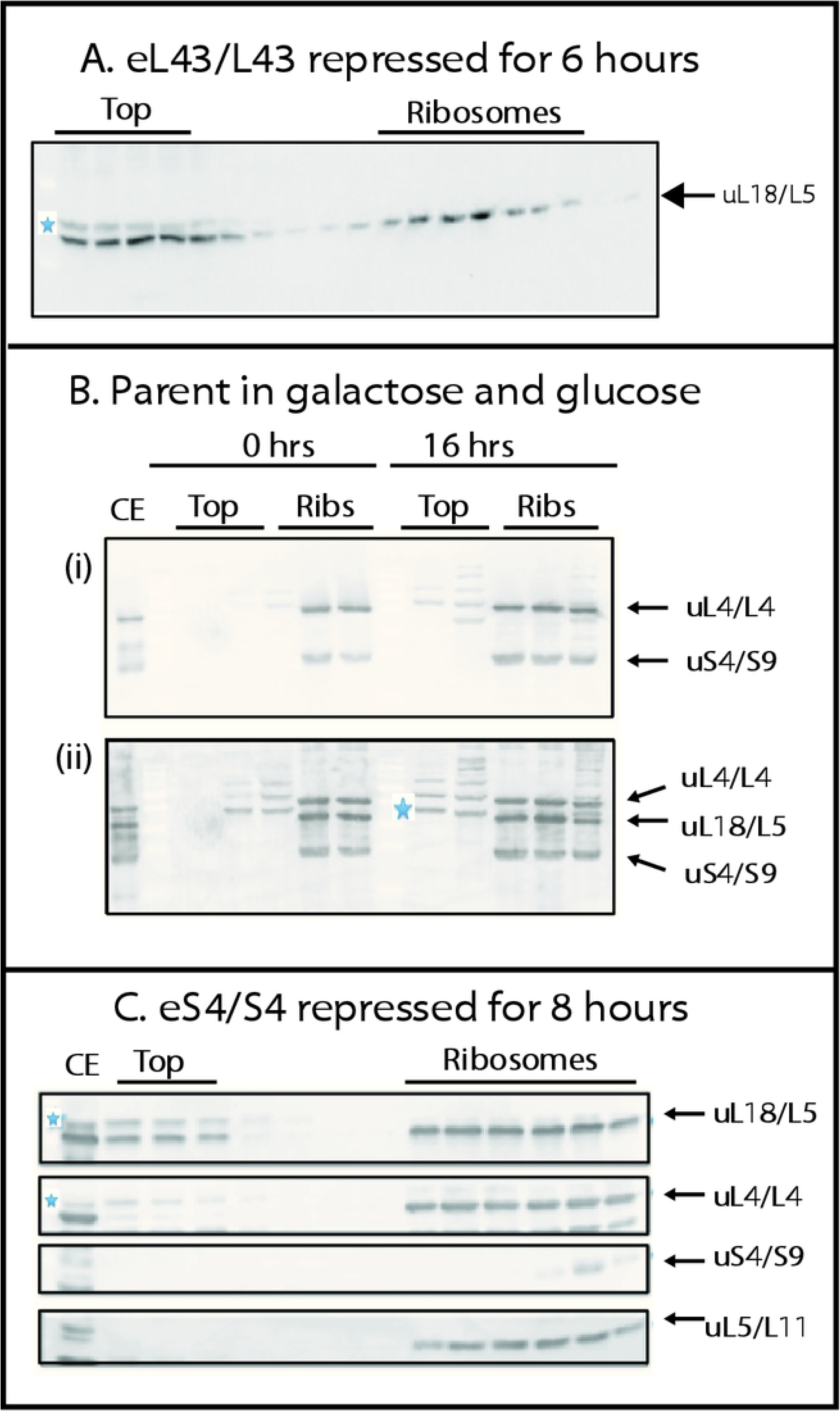
Repression of several 40S r-protein genes causes accumulation of extra-ribosomal uL18/L5, but not extra-ribosomal uL5/L11, uL4/L4, or uS4/S9. Whole cell lysates of glucose cultures of P_gal_-eL43/L43, P_gal_-eS4/S4, and the parent strain BY4741 were fractionated on sucrose gradients and the indicated fractions were analyzed by western blots probed with anti-sera for r-proteins uL18/L5, uL4/L4, eS4/S4, and uL5/L11 as indicated at each blot. (A) Pgal-eL43/L43 after 6 hours in glucose medium. (B) The parent strain (BY4741) after 0 and 16 hours in glucose. Image (i) shows a blot probed with a mixture of antisera for uL4/L4 and uS4/S9. Panel (ii) shows the same blot after it was probed further with a mixture of antisera for uL18/L5 and uL5/L11. The bands marked with a blue asterisk in some panels are not related to uL18/L5 (see Material and Methods). (C) Northern blots of fractions from a sucrose gradient loaded with P_gal_-eS4/S4 after 8 hours in glucose medium were probed with anti-sera for uS4/S9, uL4/L4, uL5/L11, and uL18/L5.

### Statistics

Pairwise t-test was used.

## Results

### Disruption of ribosome assembly

To specifically abolish assembly of 60S or 40S subunits we repressed individual r-proteins genes cognate to one or the other ribosomal subunit. This was accomplished by using yeast strains in which the only gene for a given r-protein is transcribed from the *Gal1/10* promoter. We refer to these strains as P_gal_-xx, where xx is the name of the protein encoded by the gene under galactose control. In galactose medium a full set of r-proteins is synthesized, but shifting the cells to glucose medium abrogates synthesis of r-protein xx, which prevents assembly of the cognate ribosomal subunit.

### Extra-ribosomal uL18 accumulates during repression of some 40S r-protein genes

To measure extra-ribosomal accumulation of uL18 and several other r-proteins upon repression of specific r-protein genes, we fractionated crude cell extracts on sucrose gradients and analyzed the sucrose gradient fractions on western blots probed with antisera specific to the 60S r-proteins uL18, uL5, uL4 and the 40S r-protein uS4. Fig 2A shows a western blot stained with anti-uL18 of fractions from across a sucrose gradient loaded with an extract of P_gal_-eL43 prepared 6 hours after the shift to glucose medium. A band co-migrating with the ribosomal uL18 band was observed close to the top of the sucrose gradient. A second protein marked with an asterisk and moving slightly slower also appeared. As described in Materials and Methods we confirmed that band that comigrates with the ribosomal uL18 band indeed represents uL18, while the slightly slower moving band is not related to uL18 (Fig 1). In an experiment with P_gal_-eL43 expressing FLAG-tagged uL18 a band comigrating with ribosomal FLAG-uL18 was also seen at the top of the sucrose gradient after the shift from galactose to glucose medium, but not before the shift (Fig 1A). Furthermore, no uL18 band was seen at the top of the gradient after shifting the parent strain to glucose medium (Fig 2B). Thus, the experiment in Fig 2A shows that uL18 accumulates outside ribosomal particles during repression of uL43 synthesis. This was anticipated, since repression of the 60S r-protein uL5 is known to provoke a buildup of extra-ribosomal uL18 [12].

We next asked if abolishment of 40S r-protein genes also triggered extra-ribosomal uL18 accumulation, we analyzed extracts of P_gal_-eS4 that had been shifted from galactose to glucose medium for 8 hours. Unexpectedly, we found a build-up of extra-ribosomal uL18 at the top of the sucrose gradient. In contrast, no uL4, uL5 or uS4 was found outside in the ribosome peaks (Fig 2C). Additionally, the parent strain BY4741 did not accumulate extra-ribosomal r-proteins whether grown in galactose or glucose, as expected since assembly of both subunits proceeds uninterrupted in the parent whether it grows glucose and galactose medium (Fig 2B). Overall the results in Fig 2 shows that disruption of 40S assembly can generate a pool of extra-ribosomal uL18. Extra-ribosomal accumulation of uL18 is thus not specific to interference with 60S assembly.

We then tested if repression of other 40S r-protein genes also cause formation of extra-ribosomal uL18. Sucrose gradients were loaded with whole-cell extracts after repressing the genes for uS4, eS6, uS17, eS19, or eS31. Extracts prepared after repression of the 60S genes for eL40 or eL43 were used as controls. In all cases, we found uL18 bands at the top of the gradient (Fig 3A), although the strength of the bands varied. Quantification of uL18 in the top and the ribosome fractions showed that the fraction of total uL18 found outside ribosomal subunits varied by several fold with the protein whose synthesis was abolished (Fig 3B). Repression of eS4 synthesis generated as much extra-ribosomal uL18 as did the repression of the two 60S r-protein genes. In contrast, the fraction of uL18 found in the extra-ribosomal fractions after abolishing eS31 synthesis was borderline visible. The extra-ribosomal uL18 in strains repressed for other 40S proteins was at intermediate levels.

**Fig 3.**
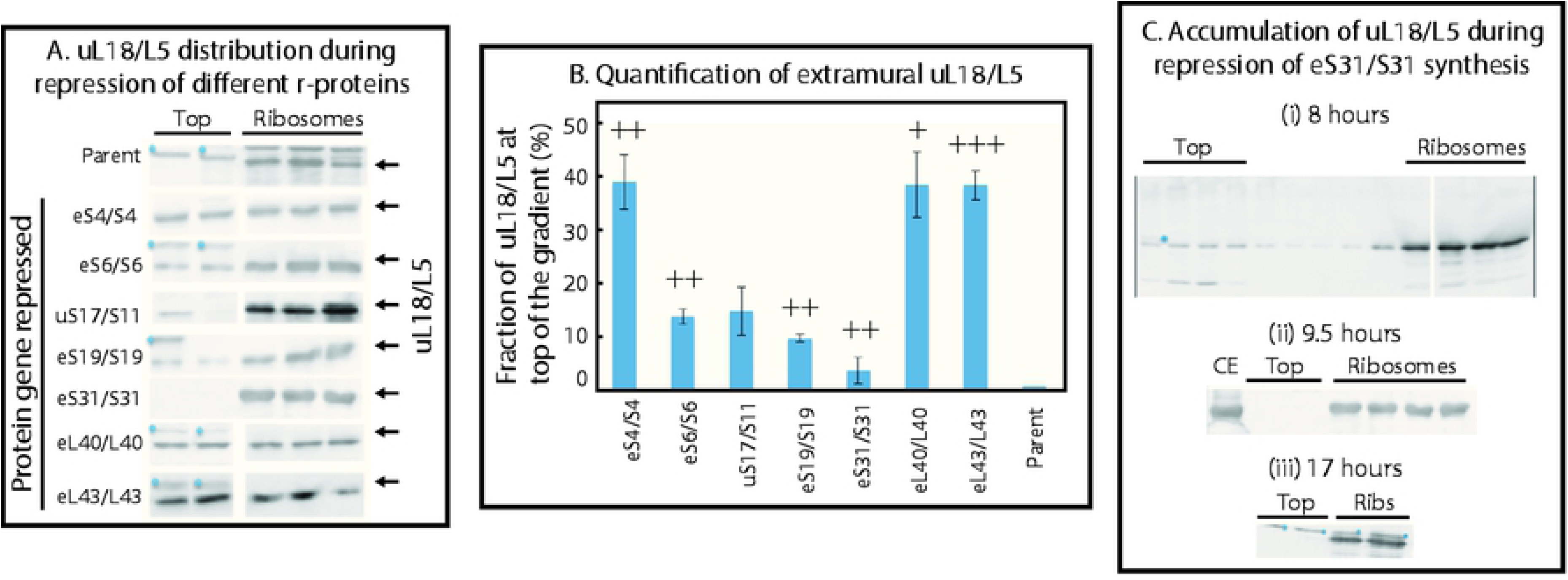
Quantification of extra-ribosomal uL18/L5 pool after repression of several 40S and 60S r-protein genes. P_gal_-eS4/S4, -eS6/S6, -uS17/S11, -eS19/S19, -eS31/S31, -eL40/L40, and -eL43/L43 were grown in galactose medium and shifted to glucose medium. (A) Sucrose gradients were loaded with lysates prepared after the shift and fractions from the top of the gradient and from the 60S-80S ribosomal peaks were analyzed on western blots probed with anti-uL18/L5. The proteins whose synthesis is repressed by the shift to glucose medium is indicated to the left of the western blots. (B) Quantification of uL18/L5 at the top of the sucrose gradient. The western blots in panel (A) were quantified using ImageJ and the fraction of the total uL18/L5 (sum of uL18/L5 in both top and ribosome fractions) present at the top of the gradient is shown. The average of three biological repeats is shown for repression eS4/S4, uS17/S11, eS31/S31, eL43/L43 genes and the average of two biological replicates is shown for the repression of the eS6/S6, eS19/S19, eL40/L40 genes. The error bars indicate standard error of the mean. The data for each gene repression experiment was compared to the results from the parent strain (BY4741) harvested after a shift to glucose medium by pairwise t-test. +++ indicates p<0.001, ++ p<0.005, + p<0.01 (C) The bands marked with a blue asterisk in some panels are not related to uL18/L5 (see Materials and Methods).

The differences among strains in the extra-ribosomal uL18 pool size led us to question whether more extra-ribosomal uL18 would accumulate with longer times of 40S r-protein gene repression. We used the Pgal-eS31 strain to address this question, because the accumulation of extra-ribosomal uL18 was barely visible in this strain. However, comparing lysates prepared after incubation in glucose medium for 8, 9.5 or 17 hours did not reveal an increase in extra-ribosomal uL18 with time, making it unlikely that the amount of extra-ribosomal uL18 changes with time (Fig 3C). Rather, the results suggest that the pool of extra-ribosomal uL18 reaches a steady-state level. Together the results in Fig 3 show that the size of the pool of extra-ribosomal uL18 during disruption of 40S assembly varies with the 40S gene repressed.

### uL18 accumulates due to interference with subunit assembly, not degradation of mature subunits

We have previously shown that the mature 40S and 60S ribosomal subunits depend on each other for stability and accumulation [9]. Thus, there are two possible principle sources of extra-ribosomal L18: modification of 60S assembly and breakdown of mature 60S subunits. To distinguish these possibilities, we investigated if blocking protein synthesis with cycloheximide changed the amount of extra-ribosomal uL18. If the extra-ribosomal uL18 stems from degradation of preexisting ribosomes, cycloheximide should not affect the pool of extra-ribosomal uL18, but if the extra-ribosomal uL18 depends on continual protein synthesis, addition of cycloheximide should reduce the pool of extra-ribosomal uL18. Accordingly, we grew P_gal_-eL43 in galactose and shifted it to glucose medium for 6 hours. At this time approximately 50% of the total uL18 was found at the top of the gradient (Fig 4A(i)). The culture was then split and cycloheximide (100 µg/ml) was added to one aliquot, while nothing was added to the other part. After 4 hours of additional culturing, both aliquots were harvested and analyzed for extra-ribosomal uL18. No uL18 band was seen at the top of the gradient after cycloheximide inhibition of protein synthesis (Fig 4A(ii)), while the level of extra-ribosomal uL18 was unchanged in the sample without the drug (Fig 4A(iii)).

**Fig 4.**
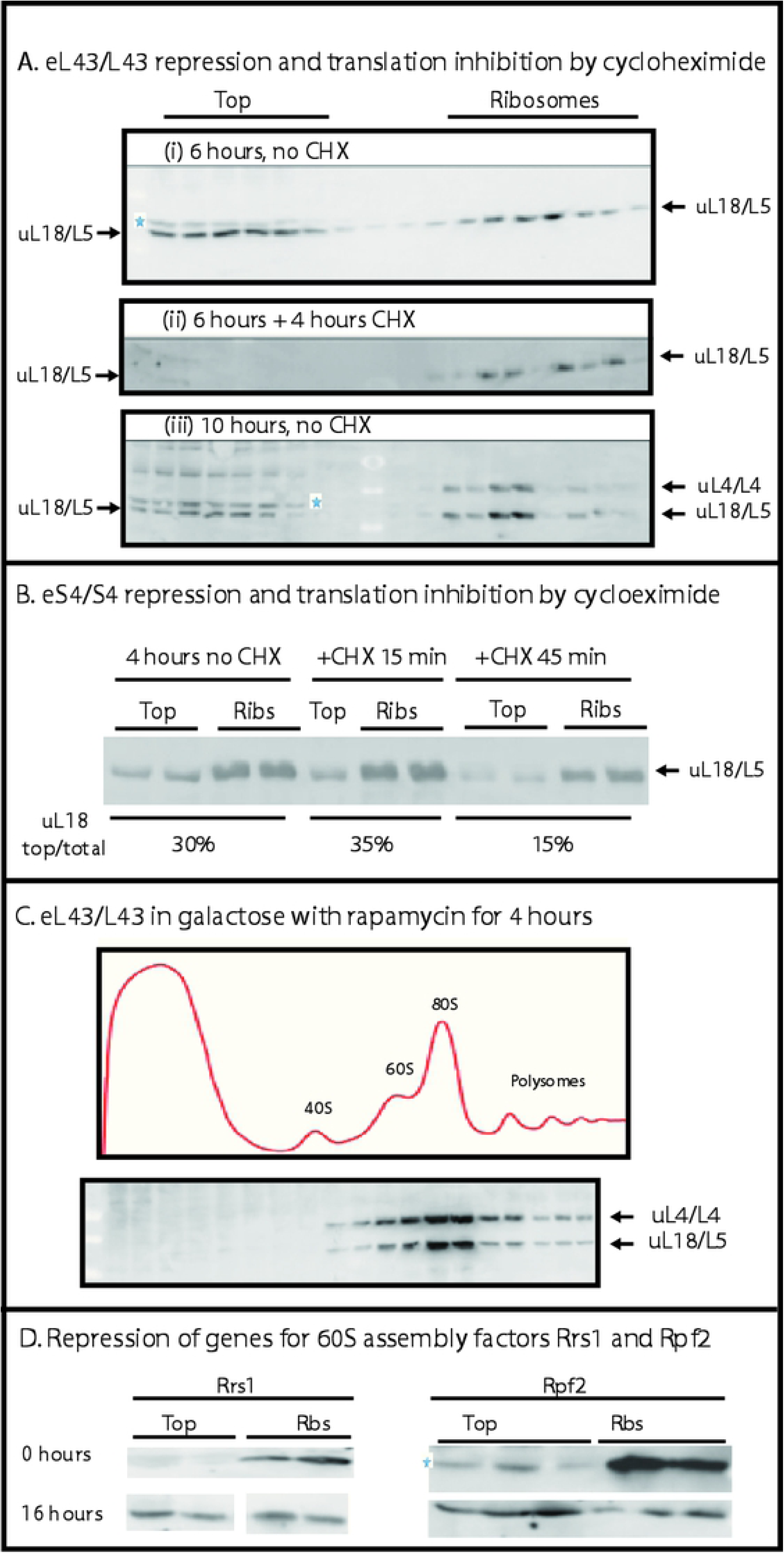
Mechanism of accumulation of extra-ribosomal uL18/L5. Cultures were grown in galactose medium and shifted to glucose medium. (A) Cycloheximide decreases the pool of extra-ribosomal uL18/L5 during repression of a 60S r-protein gene. Six hours after the shift of P_gal_-eL43/L43 from galactose to glucose medium, cycloheximide was added to an aliquot of the culture (final concentration100 µg/ml), while incubation of another aliquot was continued without the drug. Both aliquots were harvested at 10 hours after the shift from galactose to glucose. Whole cell extracts were analyzed by sucrose gradients and western blots. (i) P_gal_-eL43/L43 6 hours after the shift of media, (ii) Pgal-eL43/L43 incubated with cycloheximide added 6 hours after the media shift and harvested 10 hours after the shift. (iii) P_gal_-eL43/L43 incubated for 10 hours in glucose without cycloheximide. (B) Cycloheximide also decreases the pool of extra-ribosomal uL18/L5 during inhibition of 40S assembly. P_gal_-eS4/S4 was grown in galactose medium and shifted to glucose medium. After 4 hours in glucose medium, cycloheximide (100 µg/ml) was added and cells were harvested 0, 15 and 45 minutes after the inhibition of protein synthesis. (C) Inhibition of TOR and repression of rRNA synthesis does not result in accumulation of extra-ribosomal uL18/L5. Rapamycin was added to Pgal-eL43/L43 growing in **galactose** medium (**no** shift to glucose) and 4 hours later an extract was analyzed by sucrose gradient centrifugation and western blots developed with antisera specific to uL18/L5 and uL4/L4. (D) Repression the genes for the 60S ribosomal assembly factors Rrs1 and Rpf2 results in accumulation of extra-ribosomal uL18/L5. P_gal_-Rrs1 and -Rpf2 were grown in galactose medium and shifted to glucose medium for 16 hours. Whole cell extracts were analyzed by sucrose gradient centrifugation and western blots developed with antiserum specific to uL18/L5. The bands marked with a blue asterisk is not related to uL18/L5; see Material and Methods.

Inhibition of protein synthesis in P_gal_-eS4 gave a similar result. Cycloheximide was administrated for 15 and 45 minutes to a culture four hours after the shift from galactose to glucose. While no change was seen after 15 minutes, the extra-ribosomal uL18 level was reduced by about 50% after 45 minutes with cycloheximide (Fig 4B). Together the experiments in Fig 4A-B show that the extra-ribosomal uL18 is depleted, if it is not replenished by new synthesis, indicating that extra-ribosomal uL18 is generated during 60S assembly rather than degradation of mature 60S subunits.

To investigate if extra-ribosomal uL18 also accumulates when rRNA synthesis th is inhibited by the TOR-targeting drug rapamycin, we grew Pgal-eL43 in **galactose** medium (i.e. eL43 synthesis is **not** interrupted) and added rapamycin for 4 hours at a final concentration of 0.2 µg/ml. Fig 4C shows that no uL18 is seen at the top of the gradient after blocking rRNA transcription for 6 hours. This is consistent with the conclusion that the extra-ribosomal uL18 cannot come from degradation of mature 60S subunits, but requires continual synthesis of ribosomal components and failing assembly of ribosomal particles.

We further investigated the origin of extra-ribosomal r-proteins by depleting each of the ribosomal assembly factors Rrs1 and Rpf2 that combine with uL18, uL5, and 5S rRNA prior to docking in the precursor 60S particle [17]. Fig 4D shows that depleting either Rrs1 or Rpf2 increased the pool of extra-ribosomal uL18 in agreement with the effect of mutating the *RRS1* gene [18]. This further supports our conclusion that build-up of extra-ribosomal uL18 is caused by inhibition of ribosomal assembly rather than degradation of mature ribosomes. Furthermore, we conclude that accumulation of extra-ribosomal uL18 does not require formation of the complete uL18-uL5-5S rRNA-Rrs1-Rpf2 particle, since depletion of the assembly factors does not prevent the formation of an extra-ribosomal uL18 pool. This is also supported by the fact extra-ribosomal uL5 does not accumulate proportionally to uL18 during abrogation of eS4 synthesis (Fig 2B).

## Discussion

We have shown that the 60S r-protein uL18 accumulates extra-ribosomally during repression of several 40S r-protein genes (Figs 2 and 3). Although it was known that uL18 evades rapid degradation and accumulates outside of the ribosome during abrogation of 60S assembly [12], it was unexpected that extra-ribosomal uL18 also builds up during inhibition of 40S assembly, because repression of an r-protein gene specifically halts the assembly of their own subunit, while assembly of the other subunit continues [9]. We further showed that maintenance of the pool of extra-ribosomal uL18 requires continual protein synthesis, whether provoked by disruption of the formation of the 60S or the 40S subunit (Fig 4A-B). Thus, the extra-ribosomal r-proteins must be a product of failing or distorted ribosomal assembly rather than degradation of mature 60S subunits. This conclusion is supported by the fact that uL18 accumulates outside ribosomes in response to depletion of Rrs1 and Rpf2 (Fig 4D), both of which are involved in the incorporation of uL18 into the 60S precursor particle [17].

The fraction of uL18 found outside ribosomal subunits varies with the 40S r-protein whose synthesis is abrogated. In the extremes, abolishment of eS4synthesis generates a response similar to that seen during repression of two 60S r-protein genes, while extra-ribosomal uL18 is borderline detectable during abrogation of eS31 synthesis (Fig 3). This gradient correlates with the abundance of 40S r-proteins in the 90S ribosomal particle, an early 40S assembly intermediate [19], suggesting that preventing early steps of pre-40S precursor assembly have the strongest effect on accumulation of extra-ribosomal uL18. This can be rationalized in the context of our recent finding that the 40S r-protein eS7 and the 60S r-protein uL4 coprecipitate in immune-purifications of ribosomal precursor complexes that also contain the ITS1 sequence upstream of the cleavage site [20]. This suggests that early intermediates in the 40S and 60S subunits may interact, because they co-assemble on the emerging precursor rRNA before the pre-rRNA is cleaved between the 18S and 5.8S parts. Moreover, it is known that approximately 80% of the transcripts in rapidly growing yeast cells are cleaved between the 18S and 5.8S parts while transcription is still ongoing (“co-transcriptional rRNA processing”) [21], but the cleavage is prevented by the assembly factor Rrp5 until Domain 1 of the 60S part of the transcript is completed [22]. Since two 60S r-proteins (uL4 and uL24) bind to Domain 1, the delay of pre-rRNA cleavage until Domain 1 is synthesized is compatible with co-assembly of the early 40S and early 60S precursor particle.

We therefore posit that the simplest explanation for the buildup of extra-ribosomal uL18 during inhibition of 40S assembly is that the folding of the 60S part of the early rRNA transcript is influenced by 40S r-proteins that bind to rRNA prior to separation of the subunit moieties of the emerging rRNA transcript. Even though 40S r-proteins are not required for 60S formation, such changes to early 60S folding may affect the path used for downstream 60S assembly and thereby change the kinetics of uptake of newly synthesized 60S proteins into the precursor 60S. Ultimately, this could change the propensity for accumulation of specific extra-ribosomal 60S proteins.

While the primary function of r-proteins is as components of the ribosome, r-proteins also have important extra-ribosomal functions, at least in metazoan cells. Thus, r-proteins from both ribosomal subunits have been identified as cancer drivers [23]. The mechanism for r-protein-mediated regulation of growth and cell fate presumably involves binding of r-proteins to several regulators of growth and the progression of the cell cycle during distortion of ribosomes biogenesis (“ribosomal or nucleolar stress”) [24–27]. While these functions of extra-ribosomal proteins have been intensely investigated, little is known about the origin of the extra-ribosomal r-protein pools. Since the major features of pathways for ribosomal assembly evolutionarily conservation, we suggest that our analysis in the yeast model organism contributes to understanding the complexity how ribosome assembly impacts regulation of growth. In fact, interactions between 40S and 60S incorporation of r-proteins is likely stronger in metazoans than in yeast, because a larger fraction of pre-rRNA is cleaved into subunit-specific pieces after completion of transcription (“post-transcriptionally rRNA processing”) in metazoans than in fast-growing yeast cells. The difference between the ratio of co-transcriptional and post-transcriptional is evident from Northern blots of rRNA processing intermediates in the two types of organisms: full length rRNA precursor transcript is more prevalent relative to other processing intermediates in mammalian cells (e.g. [28]) than it is in fast-growing yeast cells (e.g. [9]).

## Acknowledgments

This work was supported by grants from the National Science Foundation, USA (0920578), The Benelein Technologies, LLC, and the University of Maryland, Baltimore County. We thank Drs. Philipp Milkereit (University of Regensburg, Germany) and John Woolford (Carnegie Mellon University, Pennsylvania, USA) for strains and plasmids. We also thank Benedikte Traasdahl for help with the manuscript.

